# The right temporoparietal junction is causally associated with embodied perspective taking

**DOI:** 10.1101/832469

**Authors:** A.K. Martin, K. Kessler, S. Cooke, J. Huang, M. Meinzer

**Affiliations:** UQ Centre for Clinical Research, University of Queensland, Brisbane, Australia; Department of Psychology, The University of Kent, Canterbury, UK; Aston Neuroscience Institute, School of Life and Health Sciences, Aston University, Birmingham, UK; Department of Neurology, University Medicine Greifswald, Greifswald, Germany

## Abstract

Several theories exist pertaining to the role of the right temporoparietal junction (rTPJ) in social cognition. A prominent theory claims the rTPJ is especially associated with embodied processes. In the present study we use high-definition transcranial direct current stimulation (HD-tDCS) to provide evidence that the rTPJ is causally associated with the embodied processes underpinning perspective taking. Eighty-eight young human adults were stratified to receive either rTPJ or dorsomedial prefrontal (dmPFC) anodal HD-tDCS in a sham-controlled, double-blind, repeated-measures design. Perspective tracking (line-of-sight) and perspective taking (embodied rotation) were assessed using a visuo-spatial perspective taking (VPT) task that required understanding *what* another person could see or *how* they see it, respectively. Embodied processing was manipulated by positioning the participant in a manner congruent or incongruent with the orientation of an avatar on the screen. As perspective taking, but not perspective tracking, is influenced by bodily posture, this allows the investigation of the specific causal role for the rTPJ in embodied processing. Crucially, anodal stimulation to the rTPJ increased the effect of bodily posture during perspective taking, whereas no such effects were identified during perspective tracking, thereby providing evidence for a causal role for the rTPJ in the embodied component of perspective taking. Stimulation to the dmPFC had no effect on perspective tracking or taking. Therefore, the present study provides support for theories postulating that the rTPJ is causally involved in embodied cognitive processing relevant to social functioning.

**Significance Statement:** The ability to understand another’s perspective is a fundamental component of social functioning. Adopting another perspective is thought to involve both embodied and non-embodied processes. The present study used high-definition transcranial direct current stimulation (HD-tDCS) and provided causal evidence that the right temporoparietal junction (rTPJ) is involved specifically in the embodied component of perspective taking. Specifically, HD-tDCS to the rTPJ, but not another hub of the social brain (dmPFC), increased the effect of body posture during perspective taking, but not tracking. This is the first causal evidence that HD-tDCS can modulate social embodied processing in a site-specific and task-specific manner.

---

Humans are fundamentally social animals. The ability to operate within large social networks requires considerable cognitive capacity, often referred to as social cognition. Recently, considerations of how the body influences cognition, especially social cognition, have grown in prominence under the theory of embodied cognition (Gallese, 2007). One social cognitive process thought to involve embodied and non-embodied processes is perspective-taking (Kessler and Rutherford, 2010; Kessler and Thomson, 2010). Specifically, perspective-taking, or imagining the world from another’s point of view, is thought to rely on the ability to “put oneself in another’s shoes” or the embodied rotation of the self into the location/orientation of another. In comparison, perspective-tracking, or understanding what another person can see, simply requires a line-of-sight judgement that does not rely on embodied processes to the same extent (Michelon and Zacks, 2006; Kessler and Rutherford, 2010). Recently, the right temporoparietal junction (rTPJ) has been suggested as a key hub for embodied processing relevant to social cognition (Wang et al., 2016; Martin et al., 2018; Martin et al., 2019). In the present study, we aimed to provide causal evidence that the rTPJ is involved in a site- and task-specific manner in embodied perspective taking. Moreover, to provide the first evidence that focal, high-definition transcranial direct current stimulation (HD-tDCS) can increase embodied processing relevant to social cognition.

The rTPJ is considered a key hub of the social brain and has been linked to higher-order processes such as theory of mind (ToM; Schurz et al., 2014). However, recent research has provided evidence for the specific cognitive processes causally associated with the rTPJ. For example, Santiesteban and colleagues (2012) found that excitatory (anodal) transcranial direct current stimulation (tDCS) to the rTPJ specifically improved the ability to inhibit non-task relevant perspectives during a visual perspective taking (VPT) task. More recently, it has also been suggested that the rTPJ has a causal role in inhibiting the self-perspective, specifically for tasks involving embodied rotation into the perspective of another person or from another location (Martin et al., 2018). Moreover, Wang and colleagues (2016) demonstrated reduced embodied processing after inhibiting the rTPJ through transcranial magnetic stimulation (TMS). The task employed by Wang et al (2016) in their Magnetoencephalography - TMS (MEG-TMS) study included both perspective tracking and taking and assessed the effects of bodily posture on response times and brain oscillations. As in previous behavioural work (Kessler and Rutherford, 2010), bodily posture was shown to affect perspective taking but not tracking and was associated with enhanced theta oscillations in rTPJ. Crucially, inhibitory TMS to the rTPJ significantly reduced the embodied response time effect for perspective taking, while a subsequent study by Gooding-Williams et al (2017) corroborated the importance of rTPJ theta oscillations during perspective taking, using repetitive TMS to entrain rTPJ at either theta or alpha frequency. Theta entrainment boosted perspective taking, and the bodily posture effect, while alpha entrainment had the opposite effect.

Another key hub of the social brain is the dorsomedial prefrontal cortex (dmPFC; Schurz et al., 2014). Previous work from our group has demonstrated that anodal HD-tDCS to the dmPFC increases the influence of the other perspective during self-perspective judgements only (Martin et al., 2017a). Crucially, although a key hub of a broader social brain network, a dissociable causal role was identified from that of the rTPJ; a role which was characterised as inhibiting the self-perspective during perspective taking (Martin et al., 2018). Therefore, stimulation of this region should not affect perspective taking or tracking when only the perspective of the other is required. It therefore offers an ideal control site to provide site-specific evidence for the role of the rTPJ in embodied perspective taking.

In the present study we employ a VPT task with embodied and non-embodied components used in previous research (Kessler and Rutherford, 2010; Kessler et al., 2014; Wang et al., 2016; Gooding-Williams et al., 2017) and used focal HD-tDCS to investigate whether the rTPJ modulates embodied processing during perspective taking in a task-specific manner as indexed by an increase in the effect of bodily posture on response times during perspective taking but not tracking. We also aim to show site-specificity by demonstrating that this effect is specific to the rTPJ and not another key hub of the social brain, the dmPFC, elucidating the role of rTPJ in embodied perspective transformations.

## METHOD

### Participants

Eighty-eight healthy young adults (46 Females; 18-36yrs, mean age= 23.27, sd= 3.69) were stratified to receive either dmPFC (N=44) or rTPJ (N=44) anodal HD-tDCS in sham-controlled, double-blind, crossover studies. Stimulation order was balanced across both sites so that half received active and half received sham stimulation during the first session. The groups were comparable on years of education, Autism Spectrum Quotient (ASQ; Baron-Cohen et al., 2001b), Hospital Anxiety and Depression Scores (HADS; Zigmond & Snaith, 1983), and across most baseline cognitive measures (see Table 1). All participants were tDCS naïve, were not currently taking psychoactive medications or substances, and had no history of neurological or severe mental health issues. All participants provided written consent and completed a safety screening questionnaire prior to the testing and were compensated for their time with a small monetary compensation. The study abided by the ethical standards as per The Declaration of Helsinki (1991; p1194). Ethical clearance was granted by The University of Queensland.

**Table 1.** Demographics and baseline cognitive performance across the rTPJ and dmPFC stimulation groups

|  | rTPJ | dmPFC | *p* |
| --- | --- | --- | --- |
|  | Mean (sd) | Mean (sd) |  |
| Age | 22.45 (3.08) | 24.09 (4.07) | 0.04 |
| ASQ | 17.68 (6.19) | 17.11 (7.09) | 0.69 |
| Years of education | 13.41 (2.17) | 14.07 (2.11) | 0.15 |
| One back lmn | 2.84 (0.09) | 2.84 (0.10) | 0.96 |
| One back acc | 1.43 (0.13) | 1.37 (0.19) | 0.08 |
| Two back lmn | 2.95 (0.10) | 2.96 (0.10) | 0.86 |
| Two back acc | 1.36 (0.14) | 1.31 (0.19) | 0.18 |
| IDN lmn | 2.73 (0.09) | 2.70 (0.08) | 0.13 |
| IDN acc | 1.45 (0.28) | 1.42 (0.20) | 0.66 |
| ISL | 27.77 (3.47) | 28.60 (2.85) | 0.23 |
| ISLR | 10.16 (1.64) | 10.51 (1.45) | 0.29 |
| CPAL err | 21.75 (27.83) | 27.30 (25.93) | 0.34 |
| SET err | 14.44 (5.19) | 18.44 (10.68) | 0.03 |
| Stroop Effect | 22.23 (6.31) | 20.47 (8.46) | 0.27 |
| Phonemic fluency | 15.23 (3.80) | 17.57 (5.00) | 0.02 |
| Semantic fluency | 24.50 (5.84) | 26.16 (8.04) | 0.27 |
| SEC | 1.13 (0.12) | 1.11 (0.16) | 0.58 |
| HADS depression | 4.05 (3.10) | 3.55 (2.83) | 0.43 |
| HADS anxiety | 7.43 (3.34) | 7.18 (3.94) | 0.75 |

ASQ= Autism Spectrum Quotient; lmn = log10 milliseconds speed of reaction for correct responses; acc = accuracy; IDN = Identification Task; ISL = International Shopping List; ISLR = International Shopping List Delayed Recall; CPAL = Continuous Paired Associates Learning; SET = Set-Switching; err = errors; SEC = Socio-emotional cognition; HADS = Hospital Anxiety and Depression Scale.

### Baseline Testing

All participants completed baseline cognitive assessment to ensure the two groups (dmPFC and rTPJ stimulation sites) were comparable and that all participants were within expected age-related norms (as in our previous studies, e.g. Martin et al., 2018). Tests included the Stroop Test, phonemic and semantic verbal fluency, completed immediately following the first stimulation session. Following the second session, participants completed a computerized cognitive battery from CogState (www.CogState.com), including the tests – international shopping test, identification test, one-back, two-back, set-switching test, continuous paired associates learning test, social-emotional cognition test, and the international shopping test-delayed recall.

Minor differences between the rTPJ and dmPFC stimulation groups were identified for age, set-switching, and phonemic fluency ability (see Table 1.). All were included as covariates and found to have no effect on stimulation response and were therefore not considered further.

### Transcranial Direct Current Stimulation

Stimulation was delivered using a one-channel direct current stimulator (DC-Stimulator Plus, NeuroConn). The anode was a small circular rubber electrode (2.5mm in diameter) and the return electrode was a concentric ring placed equidistantly around the central electrode. At the rTPJ the return electrode was slightly smaller (inner/outer diameter: 7.5/9cm) than at the dmPFC (inner/outer diameter: 9.2/11.5cm) due to the position of the right ear. Current modelling has been conducted previously (Martin et al., 2017b, 2018) and demonstrated focal delivery to the target regions. Safety has also been demonstrated (Gbadeyan et al., 2016). Electrodes were held in place with electroconductive gel (Weaver Ten20 conductive paste) and an EEG cap to ensure consistent adhesion to the skin (for details see Martin et al., 2019) of tDCS setup. The dmPFC was located 65% of the distance from FZ towards the FPz using the 10-20 EEG system. The rTPJ was located at CP6 of the EEG 10-20 system. At both stimulation sites and for both sham and active stimulation, the current ramped up to 1mA over 8 seconds and ramped down over 5 seconds. In the “sham” condition, the current was maintained at 1mA for 40 seconds whereas in the active condition the current was maintained at 1mA for 20 minutes. Researchers were blinded to the stimulation condition using the “study-mode” of the DC-Stimulator (a pre-assigned code programmed into the stimulator). Participants were also blind to the stimulation condition. To avoid carryover effects, testing sessions were at least 72 hours apart.

### Visual Perspective Taking Task

Perspective-taking and -tracking were assessed using a visual perspective taking/tracking task, employed and explained in detail in previous studies (Kessler and Rutherford, 2010; Wang et al., 2016). Briefly, on a monitor a table was presented with an avatar sat at one of six locations either 60°, 110°, or 160° from the left or right of the gaze of the participant who was seated in front of a computer screen (see Figure 1). The angular disparity was included as a manipulation of how far the participant must rotate and transform their perspective in order to take the perspective of the avatar.

**Figure 1.**
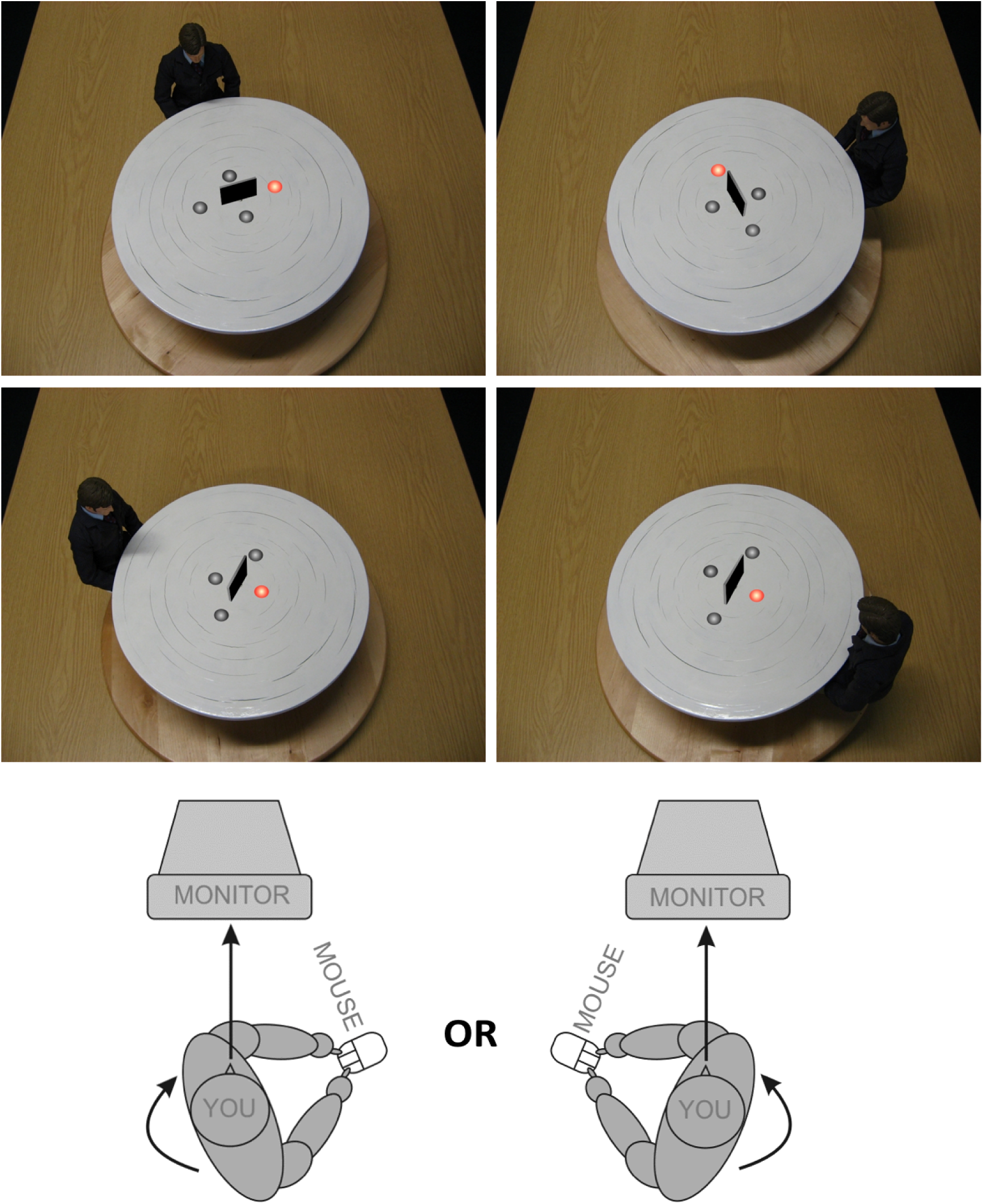
Experimental setup. The top panel displays two examples of Perspective Taking (level two VPT). Here the participant must answer whether the target (illuminated disc) is to the avatar’s left (left image) or right (right image). The middle panel displays two examples of Perspective Tracking (level one VPT). Here the participant must answer yes (right image) or no (left image) as to whether the avatar can see the illuminated disc. The bottom panel displays the body posture of the participant which was either congruent or incongruent with the avatar’s location (specifically, with the direction of mental self-rotation on any given trial). The avatar was at a disparity of either 60º, 110º, or 160º from the location of the participant. Figure adapted from Kessler & Rutherford (2010).

On the table, four grey discs were arranged around an occluding panel. On each trial, one disc would be presented in red to indicate the target. In the perspective-tracking (VPT level one) condition, participants were asked whether the disc was visible to the avatar (Yes or No response). In the perspective taking (VPT level two) condition, participants were asked whether the disc was on the avatar’s left or right (Left or Right response). In order to manipulate embodied processing, the participant’s body posture was manipulated to be either congruent or incongruent to the positioning of the avatar around the table. For instance, a body turned clockwise would be congruent with a mental rotation of the self in a clockwise direction, i.e. on trials where the avatar was seated at the left side of the table. This was achieved by asking the participant to swivel their chair to a marked position on the floor whilst maintaining their focus on the monitor. Participants were instructed to not respond until stationary. Figure 1 provides a visual representation of the experimental task setup.

Both perspective tracking and taking were presented in 14 alternating miniblocks of 24 trials each. Twelve practice trials (6 each for perspective tracking and taking) were administered at the beginning of the session to ensure participants understood task instructions

### Adverse Effects and Blinding

Adverse effects were assessed at the end of each stimulation session. Mood was assessed before and after each stimulation session (Brunoni et al., 2011) using the Visual Analogue of Mood Scale (VAMS; Folstein and Luria, 1973). Participant blinding was assessed by asking the participant “In which session do you think you received the active stimulation?” Responses could be session one or two. If a participant was not sure, they were instructed to guess.

### Experimental Design and Statistical Analysis

The Visual Perspective Taking Task was administered within a battery of social cognitive tasks that are not presented here. HD-tDCS stimulation was administered while participants were completing the Reading the Mind in the Eyes test (Baron-Cohen et al., 2001a). Following completion of this task, participants completed the VPT task followed by a task measuring socio-moral attitudes. After completion of all tasks participants completed a VAMS and baseline cognitive assessment.

All analyses were conducted in SPSS version 25. Repeated-measures Analysis of Variance (RM-ANOVA) were computed for both perspective tracking and taking conditions. The outcome was response time (RT) (for correct answers only) and the predictors were stimulation type (STIM TYPE; sham/anodal), stimulation site (STIM SITE; dmPFC/rTPJ), body posture (POSTURE; congruent/incongruent), and angle of rotation (ANGLE; 60°, 110°, 160°). Where violations of sphericity were detected, Greenhouse-Geisser corrections were employed. The task was designed to ensure accuracy was high. Therefore, we did not include analysis based on accuracy in the current study.

## RESULTS

All perspective tracking and taking response times across all conditions are presented in Table 2.

**Table 1.** Response times for perspective tracking (level one) and perspective taking (level two) during sham and anodal HD-tDCS at the rTPJ and the dmPFC.

|  | rTPJ |  | dmPFC |  |
| --- | --- | --- | --- | --- |
|  | Sham<br>Mean (sd) | Anodal<br>Mean (sd) | Sham<br>Mean (sd) | Anodal<br>Mean (sd) |
| *Perspective Tracking* |  |  |  |  |
| Congruent |  |  |  |  |
| 60° | 663.31 (188.25) | 685.25 (159.81) | 746.84 (327.31) | 751.04 (321.78) |
| 110° | 678.20 (222.92) | 705.09 (211.44) | 740.62 (293.67) | 757.18 (399.62) |
| 160° | 637.63 (185.72) | 675.79 (157.10) | 718.50 (276.32) | 762.91 (372.21) |
| Incongruent |  |  |  |  |
| 60° | 691.84 (211.56) | 698.37 (196.96) | 727.32 (310.44) | 750.39 (305.18) |
| 110° | 659.22 (192.38) | 707.16 (189.79) | 750.09 (314.41) | 748.84 (307.29) |
| 160° | 660.89 (216.83) | 662.87 (166.79) | 727.49 (289.98) | 719.38 (253.98) |
| *Perspective Taking* |  |  |  |  |
| Congruent |  |  |  |  |
| 60° | 614.79 (172.59) | 625.15 (143.94) | 663.05 (210.44) | 656.46 (162.45) |
| 110° | 670.69 (211.04) | 692.48 (192.03) | 733.01 (250.13) | 717.85 (215.70) |
| 160° | 784.00 (267.30) | 834.52 (327.83) | 837.79 (303.58) | 826.42 (285.13) |
| Incongruent |  |  |  |  |
| 60° | 636.99 (186.35) | 677.81 (182.90) | 722.42 (254.29) | 694.38 (182.17) |
| 110° | 686.08 (219.49) | 746.02 (258.74) | 801.52 (331.12) | 785.58 (344.91) |
| 160° | 814.27 (265.16) | 903.10 (421.80) | 892.38 (350.03) | 881.10 (409.45) |

**Table 2.** Adverse effects and mood change across stimulation sites and stimulation type

|  | rTPJ |  | dmPFC |  |
| --- | --- | --- | --- | --- |
|  | Sham<br>Mean (sd) | Anodal<br>Mean (sd) | Sham<br>Mean (sd) | Anodal<br>Mean (sd) |
| VAMS neg | 0.01 (1.28) | -0.28 (1.37) | -0.91 (2.04) | -1.14 (2.01) |
| VAMS pos | -0.65 (1.50) | -0.65 (2.44) | 0.30 (0.74) | 0.28 (0.83) |
| Adverse Effects | 4.16 (2.62) | 3.98 (2.66) | 4.09 (3.92) | 4.45 (3.77) |

### Perspective Tracking (Level One VPT)

As expected, bodily posture had no effect on response times, *F*(1,86)= 0.08, *p*=0.78, *η*^2^ _*p*_= 0.001. A main effect of ANGLE was identified, *F*(1.78,153.35)= 7.48, *p*=0.001, *η*^2^ _*p*_= 0.08 but the interaction between POSTURE × ANGLE was not significant, *F*(1.80,154.47)= 0.49, *p*=0.59, *η*^2^ _*p*_ = 0.01. The main effect of ANGLE was followed up with *post hoc* pairwise analysis that identified a significant difference between 160° with both 110°, *p*=0.001 and 60°, *p*=0.03. There was no difference between 110° and 60°, *p*=1.0 (All Bonferroni corrected; see Figure 2).

**Figure 2.**
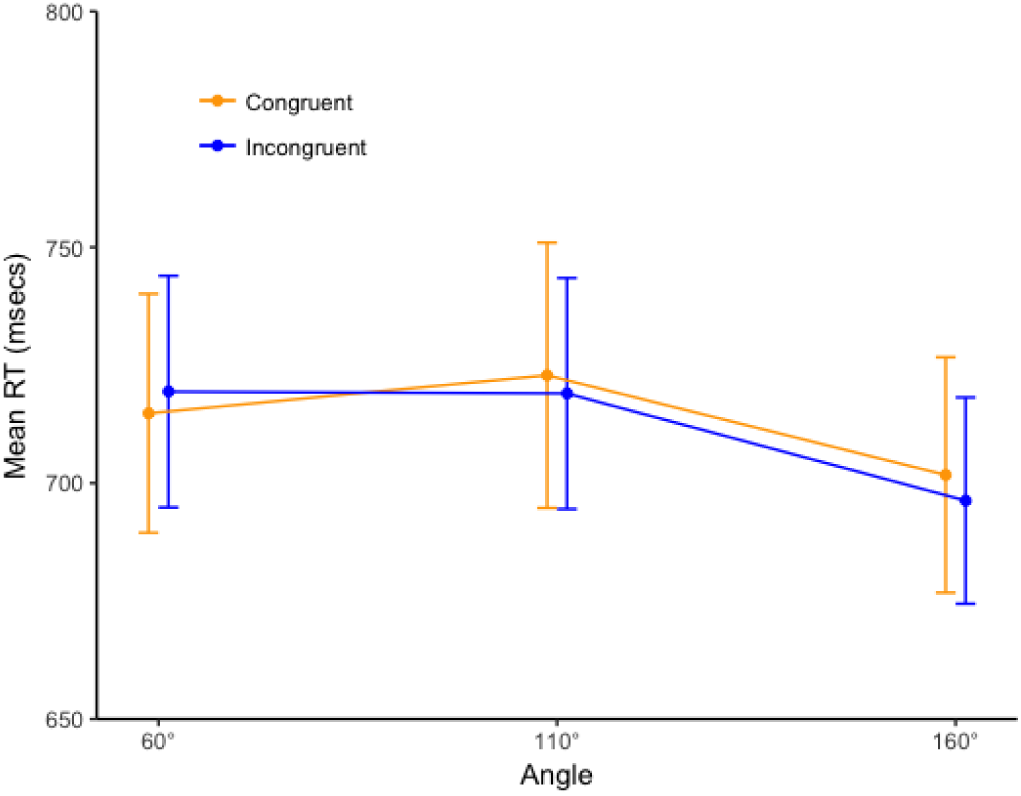
Body posture had no effect on response time for perspective tracking (Level one VPT). An effect of angle was identified such that response time was faster when the angle of difference between the participant and the avatar was 160° compared with both 60° and 110°. Stimulation to either the rTPJ or dmPFC had no effect on perspective tracking.

All stimulation effects were non-significant, STIM SITE × STIM TYPE × POSTURE × ANGLE, *F*(2, 172)= 1.29, *p*=0.28, *η*^2^ _*p*_= 0.02, STIM × POSTURE × ANGLE, *F*(2,172)= 2.60, *p*=0.08, *η*^2^ _*p*_ = 0.03, STIM × ANGLE, *F*(2,172)= 0.30, *p*=0.74, *η*^2^ _*p*_= 0.004, STIM × POSTURE, *F*(1,86)= 1.69, *p*=0.20, *η*^2^ _*p*_= 0.02, STIM TYPE × STIM SITE, *F*(1,86)= 0.10, *p*=0.80, *η*^2^ _*p*_= 0.001 and STIM TYPE, *F*(1,86)= 0.77, *p*=0.38, *η*^2^ _*p*_= 0.01. Therefore, HD-tDCS to either stimulation site did not affect perspective tracking.

### Perspective Taking (Level Two VPT)

As expected, bodily posture had a significant effect on response times, *F(*1,86)= 25.47, *p*<0.001, *η*^2^ _*p*_= 0.23 with slower responses when the participant’s bodily posture was incongruent with the location of the avatar. An effect for angle of rotation was also identified, *F(*1.32,113.59) = 84.26, *p*<0.001, *η*^2^ _*p*_ = 0.50, with response times increasing with greater angular disparity between participant and avatar. There was no interaction between the two, ANGLE × POSTURE, *F(*1.84,158.61)= 0.25, *p=*0.76, *η*^2^ _*p*_= 0.003. Therefore, angle of rotation had an effect on response time and this was comparable for congruent and incongruent bodily postures.

A significant STIM SITE × STIM TYPE × POSTURE interaction was identified, *F(*1,86)= 9.21, *p=*0.003, *η*^2^ _*p*_ = 0.10. Therefore, separate analyses were computed for the rTPJ and dmPFC stimulation sites. At the rTPJ site, a STIM TYPE × POSTURE interaction was identified, *F(*1,43)= 15.73, *p*<0.001, *η*^2^ _*p*_= 0.27. Simple effects analysed showed no significant effect of stimulation on congruent, *F(*1,43)= 0.72, *p=*0.40, *η*^2^ _*p*_= 0.02, nor on incongruent body posture, *F(*1,43)= 2.55, *p=*0.12, *η*^2^ _*p*_= 0.06. During sham stimulation, there was an effect of POSTURE, *F(*1,43)= 8.80, *p=*0.005, *η*^2^ _*p*_= 0.17. However, after anodal stimulation the effect of POSTURE was increased, *F(*1,43)= 36.44, *p*<0.001, *η*^2^ _*p*_= 0.46 (See Figure 3).

**Figure 3.**
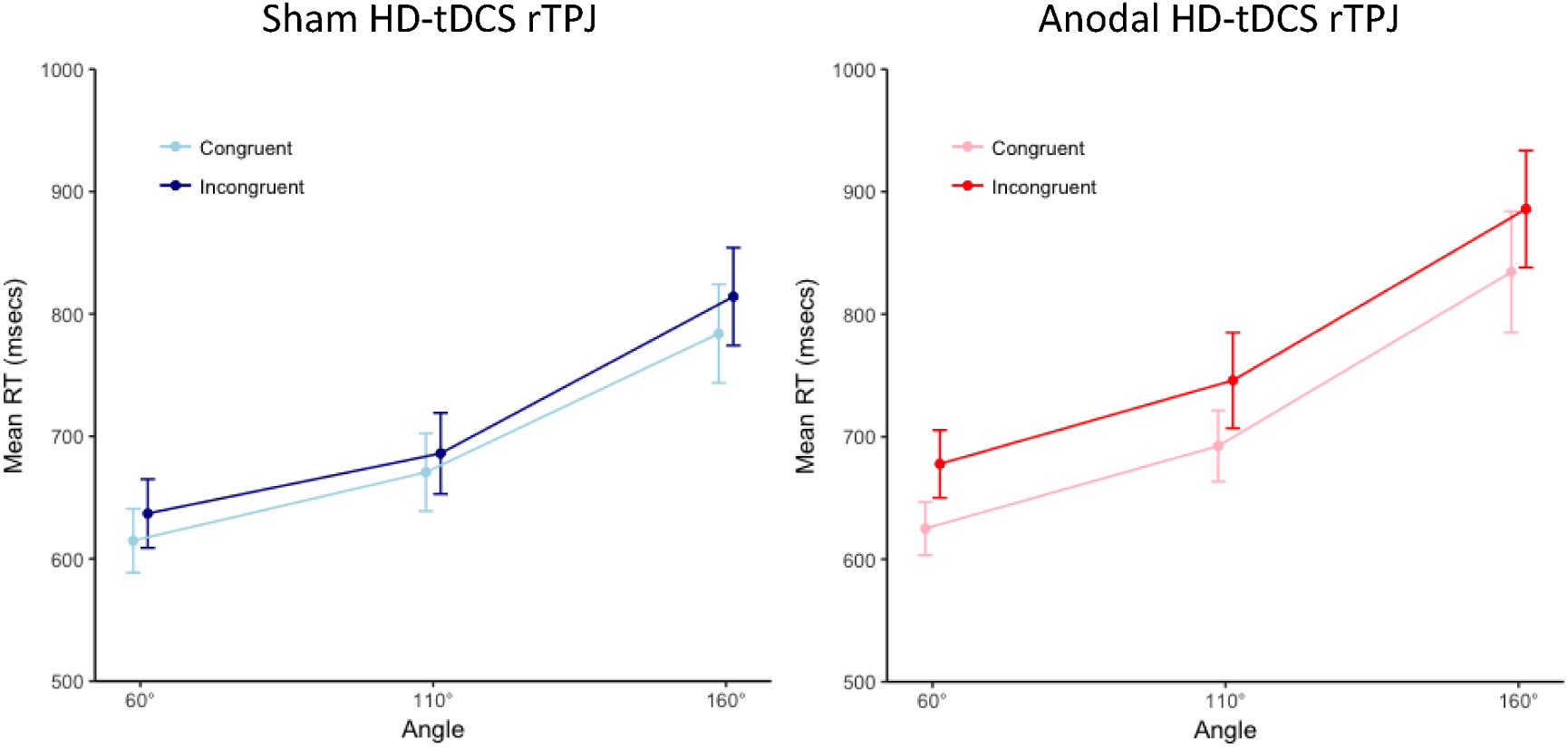
Body posture affected response times during perspective-taking (VPT level two) at baseline (Sham HD-tDCS). Anodal HD-tDCS to the rTPJ significantly increased this effect.

All stimulation effects on angle of rotation were non-significant, STIM SITE × STIM TYPE × POSTURE × ANGLE, *F(*1.78, 156.65)= 0.16, *p=*0.83, *η*^2^_*p*_ = 0.002, STIM × POSTURE × ANGLE, *F(*1.78,156.65)= 0.22, *p=*0.77, *η*^2^ _*p*_= 0.003, STIM × ANGLE, *F(*1.36,119.61)= 0.76, *p=*0.42, *η*^2^ _*p*_= 0.01.

At the dmPFC site, the STIM TYPE × POSTURE interaction was not significant, *F(*1,43)= 0.58, *p=*0.45, *η*^2^ _*p*_ = 0.01. The main effect of STIM TYPE was also not significant, *F(*1,43)= 0.31, *p=*0.58, *η*^2^ _*p*_ = 0.01.

Therefore, anodal stimulation to the rTPJ had a site-specific and task-specific effect on the embodied component of perspective taking as indexed by an increased effect of bodily posture on response times.

### Adverse Effects and Blinding

Participants were able to correctly identify the stimulation order at a rate better than chance, 56/88, *p=*0.01. However, this does not explain the results as blinding was effective at the rTPJ site, 26/44, *p=*0.23 but not at the dmPFC site, 30/44, *p=*0.02. There was no significant difference for accuracy of guessing stimulation order between the two stimulation sites, *p=*0.38. Therefore, the site- and task-specific effects are not due to a lack of participant blinding.

Stimulation had no effect on negative mood change, *F(*1,86)= 1.36, *p=*0.25 and there was no interaction with Stimulation Site, *F(*1,86)=0.24, *p=*0.88. Likewise, stimulation had no effect on positive mood change, *F(*1,86)=0.001, *p=*0.98 and there was no interaction with Stimulation Site, *F(*1,86)= 0.001, *p=*0.98. There was no difference between sham and anodal stimulation sessions for adverse effects, *F(*1,86)= 0.05, *p=*0.83 with no interaction with Stimulation Site, *F(*1,86)= 0.42, *p=*0.52. Data presented in Table 2.

## DISCUSSION

This is the first study to demonstrate that anodal HD-tDCS to the rTPJ can amplify embodied processing during visual perspective taking. Moreover, it provides site- and task-specific evidence for the efficacy of HD-tDCS to modulate specific embodied cognitive processes relevant to social functioning. The results therefore support the theory that the rTPJ is causally involved in embodied processes relevant for social cognition (Arzy et al., 2006; Wang et al., 2016; Martin et al., 2018; Martin et al., 2019). Anodal HD-tDCS to the rTPJ increased the effect of body posture on perspective taking corroborating previous evidence using TMS, which found reduced embodied processing after inhibiting the rTPJ (Wang et al., 2016), yet, faster perspective taking and enhanced embodied facilitation after entraining rTPJ at theta frequency in contrast to alpha frequency (Gooding-Williams et al., 2017).

More broadly, the rTPJ has been implicated in several aspects of self-processing including self-other distinction (Santiesteban et al., 2012, 2015; Wang et al., 2016; Payne and Tsakiris, 2017; van Elk et al., 2017), own-body imagery (Blanke and Arzy, 2005; Blanke et al., 2005), and agency (Ruby and Decety, 2001). This notion is strengthened by clinical research showing disembodiment following invasive stimulation in a patient undergoing epilepsy treatment (Blanke et al., 2002), intra-brain recordings in a patient with epilepsy linking TPJ to perspective transformations and so-called out-of-body experiences (e.g. Blanke et al., 2005; for review, Kessler and Braithwaite, 2016), as well as evidence from lesion studies (Ionta et al., 2011; Martinaud et al., 2017). Embodiment may be the key underlying process that unites the role of the rTPJ in these varied aspects of self-processing.

It is important to point out that rTPJ does not operate in isolation, but appears to be an important network hub for embodied perspective transformations, operating within a wider cortical network at theta frequency, as corroborated by recent MEG work (Bogels et al., 2015; Wang et al., 2016; Seymour et al., 2018) as well as frequency-tuned TMS entrainment (Gooding-Williams et al., 2017). Using Granger causality and imaginary coherence analysis, Seymour et al (2018) reported that rTPJ appeared to be modulated top-down at theta frequency by executive areas in the prefrontal cortex (dorsal anterior cingulate cortex and lateral prefrontal cortex) and was coupling at theta frequency with social processing areas (medial prefrontal cortex, posterior cingulate cortex) and body/action-related areas (supplementary motor area, sensorimotor cortex, posterior parietal cortex). At the same time rTPJ was desynchronising with the ventral visual stream, suggesting that rTPJ might control the switch from external events to internal states and information manipulation such as embodied mental simulations (see also Bzdok et al., 2013; Wu et al., 2015). The division of labour between TPJ and executive and social processing areas in the prefrontal cortex during embodied perspective taking has now further been corroborated by the current results, as well as by our previous anodal HD-tDCS stimulation studies (Martin et al., 2018; Martin et al., 2019). Using anodal HD-tDCS Martin et al (2017a) were able to characterise the role of dmPFC as crucial to suppressing the egocentric perspective. Since only the other’s perspective was relevant in the current task, suppression of the egocentric perspective was not required on a trial-by-trial basis and therefore dmPFC stimulation did not modulate perspective taking behaviour. The current study therefore extends previous findings to show a regionally specific effect on the distinct embodied processes underlying perspective taking ability.

Embodiment is increasingly thought to be relevant for understanding clinical conditions such as autism and psychosis (De Jaegher, 2013; Eigsti, 2013; Tschacher et al., 2017; Szczotka and Majchrowicz, 2018; Crespi and Dinsdale, 2019) and may be associated with social functioning deficits (Gallese, 2007; Goldman and de Vignemont, 2009). Moreover, older adults may use less embodied strategies (Costello and Bloesch, 2017) coupled with reduced social cognitive performance (Moran et al., 2012). Recently, non-invasive brain stimulation has shown considerable promise as a method for enhancing embodied processes (Wang et al., 2016; Lira et al., 2018; Martin et al., 2018; Hornburger et al., 2019; Martin et al., 2019) and our present and previous results across several social cognitive tasks (Martin et al., 2017a; Martin et al., 2017b, 2018; Martin et al., 2019), suggest that HD-tDCS offers an exciting new technique for improving specific social cognitive processes across a range of cohorts, especially when based on modelling of electric current flow (Martin et al., 2017b, 2018). Therefore, future research investigating the efficacy of HD-tDCS to improve embodied social cognitive processes in clinical groups is warranted.

In addition to the novel effects of anodal HD-tDCS to rTPJ, we replicate behavioural evidence for embodied processes being specific for level two perspective taking in contrast to level one perspective tracking. We further replicate previous studies regarding a slight, but consistent decrease in response latencies with increasing angular disparity for perspective tracking – which is contrary to the effect observed for perspective taking (e.g. Kessler et al, 2014; Kessler & Rutherford, 2010; Wang et al., 2016, MEG experiment). While this decrease in response times was continuous in previous studies (e.g. Wang et al., 2016, MEG experiment), here we observed a more discontinuous pattern with a significant drop-off only at 160 deg, which might be linked to a clearer dissociation between self and other perspective at high angular disparities (see Kessler et al., 2014, for a detailed discussion).

Despite consistent behavioural evidence for the efficacy of tDCS to affect social cognitive processes (Sellaro et al., 2016), little is known about how tDCS affects brain function. However, recent evidence suggests that HD-tDCS to the rTPJ increases low-frequency oscillatory activity that may exert inhibitory effects at the network-level and enable switching between endogenous and exogenous processing streams (Donaldson et al., 2019). Further research is required combining HD-tDCS and EEG during social cognitive tasks to investigate how electrical stimulation interacts with intrinsic neural processes. The HD-tDCS set-up used in the present study is compatible with the MRI environment (Gbadeyan et al., 2016) which should motivate future research into how HD-tDCS to social brain regions such as the rTPJ affects neural functioning locally at the stimulation site and at more distant but functionally connected regions within a broader network. Local and network level effects of conventional tDCS have been demonstrated (Keeser et al., 2011; Stagg and Nitsche, 2011; Meinzer et al., 2012). Understanding the systems level effect of HD-tDCS will improve our mechanistic understanding of how tDCS affects the brain in a physiologically relevant manner. Such research will complement the ongoing research providing neurophysiological evidence for the efficacy of tDCS to affect brain function (Huang et al., 2018). Replicated, task-specific, and site-specific evidence such as the evidence for a causal role for the rTPJ in embodied perspective taking from the present study and others (Martin et al., 2018; Martin et al., 2019) increase the evidence for HD-tDCS as a valid scientific technique with potential for clinical applications.

## CONCLUSION

Anodal HD-tDCS to the rTPJ, but not to the dmPFC, increased the effect of body posture during perspective taking, but not during perspective tracking, thereby providing the first causal evidence that HD-tDCS can modulate social embodied processing in a site-specific and task-specific manner.

## Notes

#### Summary of Updates

Updated abstract to match the manuscript

## REFERENCES

Arzy S, Thut G, Mohr C, Michel CM, Blanke O (2006) Neural basis of embodiment: distinct contributions of temporoparietal junction and extrastriate body area. J Neurosci 26:8074–8081.

Baron-Cohen S, Wheelwright S, Hill J, Raste Y, Plumb I (2001a) The “Reading the Mind in the Eyes” Test revised version: a study with normal adults, and adults with Asperger syndrome or high-functioning autism. J Child Psychol Psychiatry 42:241–251.

Baron-Cohen S, Wheelwright S, Skinner, R, Martin, J, Clubley, E (2001b) The autism-spectrum quotient (AQ): evidence from Asperger syndrome/high-functioning autism, males and females, scientists and mathematicians. J Autism Dev Disord 31:5–17

Blanke O, Arzy S (2005) The out-of-body experience: disturbed self-processing at the temporo-parietal junction. Neuroscientist 11:16–24.

Blanke O, Ortigue S, Landis T, Seeck M (2002) Stimulating illusory own-body perceptions. Nature 419:269–270.

Blanke O, Mohr C, Michel CM, Pascual-Leone A, Brugger P, Seeck M, Landis T, Thut G (2005) Linking out-of-body experience and self processing to mental own-body imagery at the temporoparietal junction. J Neurosci 25:550–557.

Bogels S, Barr DJ, Garrod S, Kessler K (2015) Conversational Interaction in the Scanner: Mentalizing during Language Processing as Revealed by MEG. Cereb Cortex 25:3219–3234.

Brunoni AR, Amadera J, Berbel B, Volz MS, Rizzerio BG, Fregni F (2011) A systematic review on reporting and assessment of adverse effects associated with transcranial direct current stimulation. Int J Neuropsychopharmacol 14:1133–1145.

Bzdok D, Langner R, Schilbach L, Jakobs O, Roski C, Caspers S, Laird AR, Fox PT, Zilles K, Eickhoff SB (2013) Characterization of the temporo-parietal junction by combining data-driven parcellation, complementary connectivity analyses, and functional decoding. NeuroImage 81:381–392.

Costello MC, Bloesch EK (2017) Are Older Adults Less Embodied? A Review of Age Effects through the Lens of Embodied Cognition. Front Psychol 8:267.

Crespi B, Dinsdale N (2019) Autism and psychosis as diametrical disorders of embodiment. Evol Med Public Health 2019:121–138.

De Jaegher H (2013) Embodiment and sense-making in autism. Front Integr Neurosci 7:15.

Donaldson PH, Kirkovski M, Yang JS, Bekkali S, Enticott PG (2019) High-definition tDCS to the right temporoparietal junction modulates slow-wave resting state power and coherence in healthy adults. J Neurophysiol 122:1735–1744.

Eigsti IM (2013) A review of embodiment in autism spectrum disorders. Front Psychol 4:224.

Folstein MF, Luria R (1973) Reliability, validity, and clinical application of the Visual Analogue Mood Scale. Psychol Med 3:479–486.

Gallese V (2007) Before and below ‘theory of mind’: embodied simulation and the neural correlates of social cognition. Philos Trans R Soc Lond B Biol Sci 362:659–669.

Gbadeyan O, Steinhauser M, McMahon K, Meinzer M (2016) Safety, Tolerability, Blinding Efficacy and Behavioural Effects of a Novel MRI-Compatible, High-Definition tDCS Set-Up. Brain Stimul 9:545–552.

Goldman A, de Vignemont F (2009) Is social cognition embodied? Trends Cogn Sci 13:154–159.

Gooding-Williams G, Wang H, Kessler K (2017) THETA-Rhythm Makes the World Go Round: Dissociative Effects of TMS Theta Versus Alpha Entrainment of Right pTPJ on Embodied Perspective Transformations. Brain Topogr 30:561–564.

Hornburger H, Nguemeni C, Odorfer T, Zeller D (2019) Modulation of the rubber hand illusion by transcranial direct current stimulation over the contralateral somatosensory cortex. Neuropsychologia 131:353–359.

Huang Y, Liu AA, Lafon B, Friedman D, Dayan M, Wang X, Bikson M, Doyle WK, Devinsky O, Parra LC (2018) Correction: Measurements and models of electric fields in the in vivo human brain during transcranial electric stimulation. Elife 7.

Ionta S, Heydrich L, Lenggenhager B, Mouthon M, Fornari E, Chapuis D, Gassert R, Blanke O (2011) Multisensory mechanisms in temporo-parietal cortex support self-location and first-person perspective. Neuron 70:363–374.

Keeser D, Meindl T, Bor J, Palm U, Pogarell O, Mulert C, Brunelin J, Moller HJ, Reiser M, Padberg F (2011) Prefrontal transcranial direct current stimulation changes connectivity of resting-state networks during fMRI. J Neurosci 31:15284–15293.

Kessler K, Rutherford H (2010) The Two Forms of Visuo-Spatial Perspective Taking are Differently Embodied and Subserve Different Spatial Prepositions. Front Psychol 1:213.

Kessler K, Thomson LA (2010) The embodied nature of spatial perspective taking: embodied transformation versus sensorimotor interference. Cognition 114:72–88.

Kessler K, Cao L, O’Shea KJ, Wang H (2014) A cross-culture, cross-gender comparison of perspective taking mechanisms. Proc Biol Sci 281:20140388.

Lira M, Pantaleao FN, de Souza Ramos CG, Boggio PS (2018) Anodal transcranial direct current stimulation over the posterior parietal cortex reduces the onset time to the rubber hand illusion and increases the body ownership. Exp Brain Res 236:2935–2943.

Martin AK, Su P, Meinzer M (2019) Common and unique effects of HD-tDCS to the social brain across cultural groups. Neuropsychologia 133:107170.

Martin AK, Dzafic I, Ramdave S, Meinzer M (2017a) Causal evidence for task-specific involvement of the dorsomedial prefrontal cortex in human social cognition. Soc Cogn Affect Neurosci.

Martin AK, Huang J, Hunold A, Meinzer M (2017b) Sex Mediates the Effects of High-Definition Transcranial Direct Current Stimulation on “Mind-Reading". Neuroscience 366:84–94.

Martin AK, Huang J, Hunold A, Meinzer M (2018) Dissociable roles within the social brain for self other processing: a HD-tDCS study. Cereb Cortex in press.

Martinaud O, Besharati S, Jenkinson PM, Fotopoulou A (2017) Ownership illusions in patients with body delusions: Different neural profiles of visual capture and disownership. Cortex 87:174–185.

Meinzer M, Antonenko D, Lindenberg R, Hetzer S, Ulm L, Avirame K, Flaisch T, Floel A (2012) Electrical brain stimulation improves cognitive performance by modulating functional connectivity and task-specific activation. J Neurosci 32:1859–1866.

Michelon P, Zacks JM (2006) Two kinds of visual perspective taking. Percept Psychophys 68:327–337.

Moran JM, Jolly E, Mitchell JP (2012) Social-cognitive deficits in normal aging. J Neurosci 32:5553–5561.

Payne S, Tsakiris M (2017) Anodal transcranial direct current stimulation of right temporoparietal area inhibits self-recognition. Cogn Affect Behav Neurosci 17:1–8.

Ruby P, Decety J (2001) Effect of subjective perspective taking during simulation of action: a PET investigation of agency. Nat Neurosci 4:546–550.

Santiesteban I, Banissy MJ, Catmur C, Bird G (2012) Enhancing social ability by stimulating right temporoparietal junction. Curr Biol 22:2274–2277.

Santiesteban I, Banissy MJ, Catmur C, Bird G (2015) Functional lateralization of temporoparietal junction - imitation inhibition, visual perspective-taking and theory of mind. Eur J Neurosci.

Schurz M, Radua J, Aichhorn M, Richlan F, Perner J (2014) Fractionating theory of mind: a meta-analysis of functional brain imaging studies. Neurosci Biobehav Rev 42:9–34.

Sellaro R, Nitsche MA, Colzato LS (2016) The stimulated social brain: effects of transcranial direct current stimulation on social cognition. Ann N Y Acad Sci 1369:218–239.

Seymour RA, Wang H, Rippon G, Kessler K (2018) Oscillatory networks of high-level mental alignment: A perspective-taking MEG study. NeuroImage 177:98–107.

Stagg CJ, Nitsche MA (2011) Physiological basis of transcranial direct current stimulation. Neuroscientist 17:37–53.

Szczotka J, Majchrowicz B (2018) Schizophrenia as a disorder of embodied self. Psychiatr Pol 52:199–215.

Tschacher W, Giersch A, Friston K (2017) Embodiment and Schizophrenia: A Review of Implications and Applications. Schizophr Bull 43:745–753.

van Elk M, Duizer M, Sligte I, van Schie H (2017) Transcranial direct current stimulation of the right temporoparietal junction impairs third-person perspective taking. Cogn Affect Behav Neurosci 17:9–23.

Wang H, Callaghan E, Gooding-Williams G, McAllister C, Kessler K (2016) Rhythm makes the world go round: An MEG-TMS study on the role of right TPJ theta oscillations in embodied perspective taking. Cortex 75:68–81.

Wu Q, Chang CF, Xi S, Huang IW, Liu Z, Juan CH, Wu Y, Fan J (2015) A critical role of temporoparietal junction in the integration of top-down and bottom-up attentional control. Hum Brain Mapp 36:4317–4333.

Zigmond AS, Snaith RP (1983) The hospital anxiety and depression scale. Acta Psychiatr Scand 67: 361–370.

